# Androgen-regulated transcription of *ESRP2* drives alternative splicing patterns in prostate cancer

**DOI:** 10.1101/629618

**Authors:** Jennifer Munkley, Li Ling, S R Gokul Krishnan, Gerald Hysenaj, Emma Scott, Htoo Zarni Oo, Teresa M. Maia, Kat Cheung, Ingrid Ehrmann, Karen E. Livermore, Hanna Zielinska, Oliver Thompson, Bridget Knight, Paul McCullagh, John McGrath, Malcolm Crundwell, Lorna W. Harries, Mads Daugaard, Simon Cockell, Nuno L. Barbosa-Morais, Sebastian Oltean, David J Elliott

## Abstract

Prostate is the most frequent cancer in men. Prostate cancer progression is driven by androgen steroid hormones, and delayed by androgen deprivation therapy (ADT). Androgens control transcription by stimulating androgen receptor (AR) activity, yet also control pre-mRNA splicing through less clear mechanisms. Here we find androgens regulate splicing through AR-mediated transcriptional control of the epithelial-specific splicing regulator *ESRP2*. Both *ESRP2* and its close paralog *ESRP1* are highly expressed in primary prostate cancer. Androgen stimulation induces splicing switches in many endogenous ESRP2-controlled mRNA isoforms, including a key splicing switch in the metastatic regulator *FLNB* which is associated with disease relapse. *ESRP2* expression in clinical prostate cancer is repressed by ADT, which may thus inadvertently dampen epithelial splice programmes. Supporting this, *FLNB* splicing was reciprocally switched by the AR antagonist bicalutamide (Casodex^®^). Our data reveal a new mechanism of splicing control in prostate cancer with important implications for metastatic disease progression.

**Key points:**

- Transcriptional regulation of ESRP2 by the androgen receptor controls splice isoform patterns in prostate cancer cells.
- Splicing switches regulated by the androgen-ESRP2 axis include a splice isoform in the *FLNB* gene that is a known metastatic driver.
- Both ESRP1 and ESRP2 are highly expressed in prostate cancer tissue.
- Ectopic expression of ESRP1 and 2 inhibits prostate cancer cell growth.
- By repressing ESRP2 expression androgen deprivation therapy (ADT) may dampen epithelial splicing programmes to inadvertently prime disease progression towards metastasis.

## Introduction

Prostate is the most common male gender-specific cancer (1). Prostate cancer progression is controlled by androgen steroid hormones including testosterone and its active metabolite 5-α dihydroxytestosterone. Androgens stimulate androgen receptor (AR) signalling in prostate cancer cells to control transcription, including of genes that regulate the cell cycle, central metabolism and biosynthesis, as well as housekeeping functions (2-5). The roles of both androgens and the AR in transcription have been intensively investigated. However, androgens and the AR also regulate alternative pre-mRNA splicing through still largely unknown mechanisms (6-11). This represents a very important knowledge gap: alternative splicing patterns in cancer cells can generate protein isoforms with different biological functions (12), and is a key process in the generation of biological heterogeneity in prostate cancer (13, 14).

Androgens are also closely linked to prostate cancer treatment, with androgen deprivation therapy (ADT) being the principal pharmacological strategy for locally advanced and metastatic disease. ADT utilises drugs to inhibit gonadal and extragonadal androgen biosynthesis and additionally competitive AR antagonists block androgen binding and abrogate AR function (4). ADT delays disease progression, but after 2-3 years tumours often grow again developing castration resistance with a median survival time of 16 months (15). The central role of androgens and the AR in prostate cancer, and the poor clinical outlook of castration-resistance prostate cancer (CRPCa), have made it crucially important to identify androgen-regulated target genes and mechanisms of function –particularly those that relate to metastasis. The process of epithelial-mesenchymal transition (EMT) plays a pivotal role in prostate cancer metastasis (16-20). While the mechanisms driving EMT in prostate cancer are poorly understood, ADT has recently been shown to directly induce EMT in both mouse and human prostate tissue (21, 22). Importantly, changes in alternative splicing patterns can have dramatic effects on EMT and on metastatic disease progression (23).

While the mechanisms through which androgens regulate splicing control are not well understood, splicing itself takes place in the spliceosome, which is a multi-component structure containing a core of essential proteins and small nuclear RNAs (24). Splicing inclusion of alternative exons is often controlled by splicing regulator proteins that bind either to regulated exons or within their adjacent flanking intron sequences (25). The estrogen and progesterone steroid nuclear hormone receptors control splicing via recruitment of alternative splicing regulators (including the RNA helicases Ddx5 and Ddx17) (7, 8, 26), and by changing RNA polymerase II extension rates and chromatin structure to affect splice site selection (27, 28). Steroid hormones can also drive selection of alternative promoters to include different upstream exons in mRNAs (9, 10). However, to what extent the above mechanisms may contribute to androgen-mediated splicing is largely unknown.

We reasoned that a potential model to unify the role of androgens and the AR in transcription and splicing control could be via transcriptional regulation of genes that encode splicing regulatory proteins. Using a recently described set of genes that reciprocally change expression in response to androgen stimulation in culture and ADT in patients (29), here we identify AR-mediated transcriptional control of the key splicing regulator protein Epithelial Splicing Regulator Protein 2 (ESRP2). Importantly, many ESRP2-regulated exons switch splicing in response to androgen stimulation. ESRP2 and its close relative ESRP1 (60% identical to ESRP2 protein) are important regulators of epithelial alternative splicing patterns (12, 30-35), reduced expression of which can drive critical aspects of EMT (23, 30, 36). Our data identify an AR-ESRP2 axis controlling splicing patterns in prostate cancer cells, and further suggest that reduced ESRP2 levels in response to ADT may inadvertently prime prostate cancer cells to facilitate longer term disease progression.

## Results

### *ESRP2* is a direct target for AR regulation in prostate cancer cells

To first gain insight into how androgens may mediate patterns of splicing control we analysed a recently generated dataset of genes that have reciprocal expression patterns on acute androgen stimulation *in vitro* versus clinical ADT (29). While a number of genes encoding splicing factors changed expression in response to acute androgen stimulation *in vitro*, only one such gene *ESRP2* also showed a reciprocal expression switch between acute androgen stimulation in culture and ADT in patients (29). *ESRP2* expression decreased following ADT in 7/7 prostate cancer patients (37) (Figure 1A). Furthermore, RNAseq data from LTL331 patient-derived xenografts (38) also showed reduced *ESRP2* mRNA levels following castration (Figure 1B). These data support *in vivo* androgen-regulation of *ESRP2* transcription.

**Figure 1.**
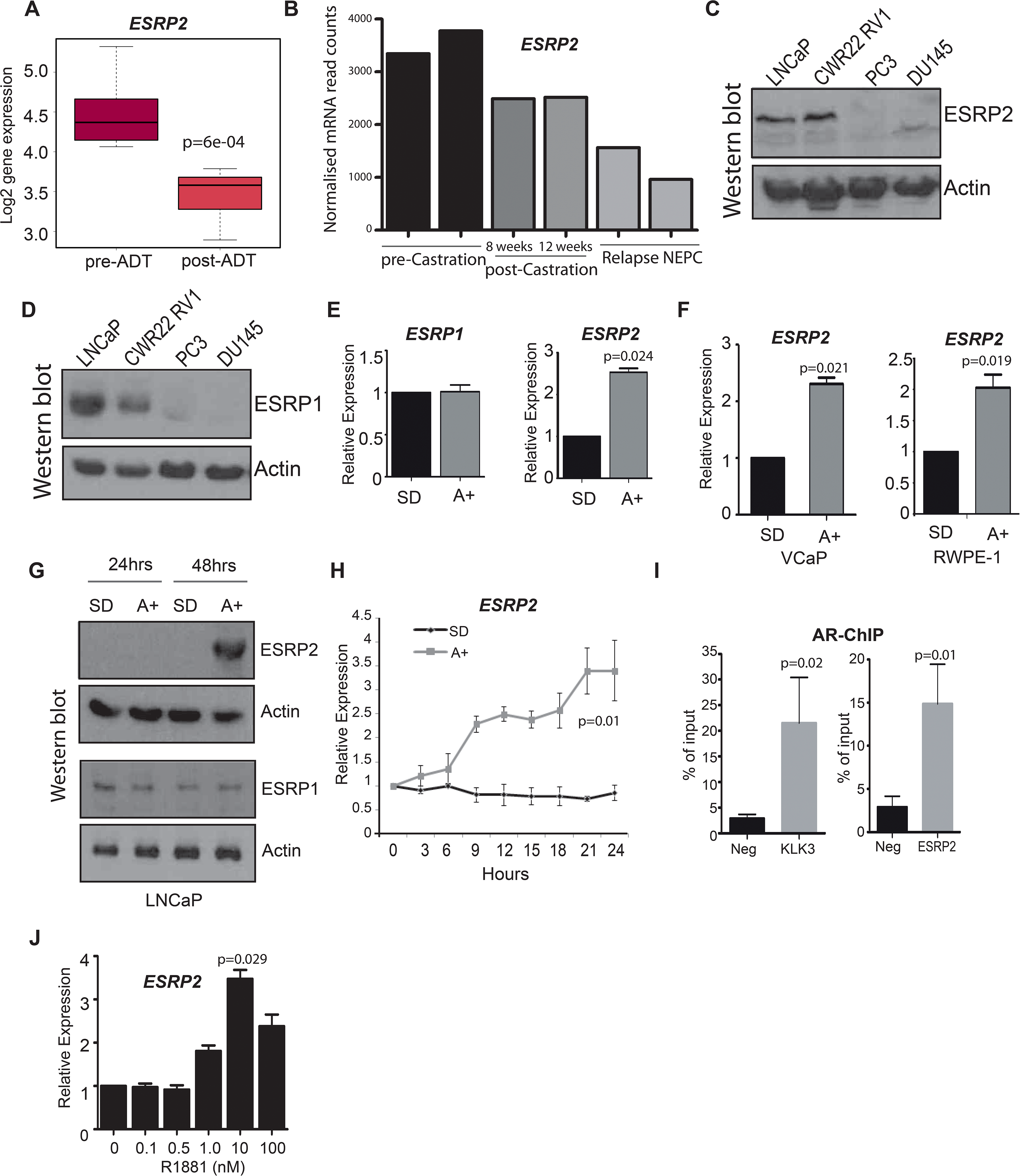
*ESRP2* is a direct target for AR regulation in prostate cancer cells. **(A)** Analysis of RNAseq data from human prostate cancer pre-and post-androgen deprivation therapy (ADT) (37, 52) shows that there is a significant downregulation of *ESRP2* mRNA following ADT in all 7 patients tested (p=6e-04, Mann Whitney U test). **(B)** RNAseq data from LTL331 patient-derived xenografts grown in mice (38) show reduced *ESRP2* mRNA levels following castration. **(C)** Western blot analysis of ESRP2 levels in a range of prostate cancer cell lines (actin was used as a loading control). **(D)** Western blot analysis of ESRP1 levels in prostate cancer cell lines. **(E)** Real-time PCR analysis of *ESRP2* and *ESRP1* mRNAs in LNCaP cells grown in steroid deplete (SD) or androgen (A+) treated conditions for 24 hours (statistical significance calculated by t test). **(F)** Real-time PCR analysis of ESRP2 mRNA in RWPE-1 cells grown in steroid deplete (SD) or androgen (A+) treated conditions for 24 hours. **(G)** Western blots analysis of ESRP1 and 2 protein in LNCaP cells treated with 10nm R1881 (androgens) for 24 and 48 hours. **(H)** Quantitative analysis (real-time PCR) of *ESRP2* mRNA expression over a 24 hour time course following androgen exposure. **(I)** Real-time PCR analysis of AR-ChIP performed in LNCaP cells treated with 10nM R1881 for 24 hours revealed AR binding proximal to the *ESRP2* gene. **(J)** Induction of *ESRP2* is evident in LNCaP cells treated with R1881 concentrations between 1 to 100 nM. Statistical significances were calculated by t tests, apart from (A) which used a Mann Whitney U test, and H which used Two-way ANOVA. Real time PCR analyses used at least 3 independent biological replicates (RNA prepared from separate samples), apart from the AR ChIP (panel I) for which each value shown is a mean of 3 technical replicates.

Further analysis of ESRP2 expression patterns in prostate cancer cell lines revealed that *ESRP2* is controlled by androgens, but not its close paralog *ESRP1*. *ESRP1* gene expression did not significantly change following castration in the LTL331 patient-derived xenografts (38). Western blots detected high endogenous levels of both ESRP1 and ESRP2 levels within the AR positive LNCaP and CWR22 RV1 prostate cancer cell lines, as compared to the AR negative PC3 and DU145 prostate cancer cell lines (Figure 1C and 1D). qPCR analysis showed that *ESRP2* gene expression in the AR-positive LNCaP cell line was activated in response to androgens, but no androgen effect was observed for *ESRP1* gene expression (Figure 1E). Androgen mediated-control of *ESRP2* expression was also detected in two additional AR-expressing prostate cell lines VCaP and RWPE-1 (Figure 1F). Induced ESRP2 protein expression was detected 48 hours after androgen exposure, with ESRP1 protein levels not changing over this same time period (Figure 1G). The specificity of the ESRP1 and ESRP2 antibodies used in these experiments was confirmed by detection of over-expressed protein and detection of siRNA mediated protein depletion by western blot (Figure 1-Figure supplement 1A and 1B).

Further experimental analysis also support *ESRP2* as an early and so likely direct target for transcriptional control by the AR: (i) *ESRP2* gene expression in LNCaP cells was rapidly induced in response to 10nM of the synthetic androgen analogue R1881 (methytrienolone) (Figure 1H). (ii) Chromatin immunoprecipitation (ChIP) from LNCaP cells confirmed direct AR binding to a site within 20Kb of the *ESRP2* gene promoter that had been previously predicted from a genome-wide study (at position chr16: 68210834-68211293 on human genome assembly HG38) (2) (Figure 1I). The AR ChIP signal adjacent to *ESRP2* was similar to that detected in parallel for *KLK3* (PSA), which is a known AR-regulated gene. (iii) Consistent with *ESRP2* regulation at physiological androgen concentrations, *ESRP2* transcription in LNCaP cells was induced over a wide range of R1881 concentrations ranging from 1 nM to 100 nM (Figure 1J). Each of these above data are consistent with AR - mediated regulation of *ESRP2* expression levels.

### *ESRP2* and its paralog *ESRP1* are highly expressed in primary prostate tumours and inhibit tumour growth *in vivo*

We next monitored *ESRP1* and *ESRP2* expression profiles from prostate cancer patients. Meta-analysis of 719 clinical prostate cancer tumours from 11 previously published studies detected significant up-regulation of both *ESRP1* and *ESRP2* in 9/11 datasets (Figure 2-source data 1) (39-50). We experimentally validated this meta-analysis using two independent panels of clinical samples. Real-time PCR showed significant up-regulation of both *ESRP1* and *ESRP2* mRNA in (1) prostate carcinoma relative to benign prostate hyperplasia (BPH) (Figure 2A); and (2) in 9 prostate tumour samples relative to matched normal tissue from the same patient (Figure 2B). A recent study by Walker et al (2017) identified a molecular subgroup of prostate cancers with metastatic potential at presentation (51). Within this dataset *ESRP1* was 2.76 fold up-regulated in the ‘metastatic-subgroup’ compared to the ‘non-metastatic subgroup’. Using RNA from a subset of samples from the Walker et al. study, we confirmed significant (p < 0.05) upregulation of the *ESRP1* gene in primary prostate cancer patients presenting with a metastatic biology (Figure 2C). *ESRP2* gene expression did not significantly increase in the 20 samples studied.

**Figure 2.**
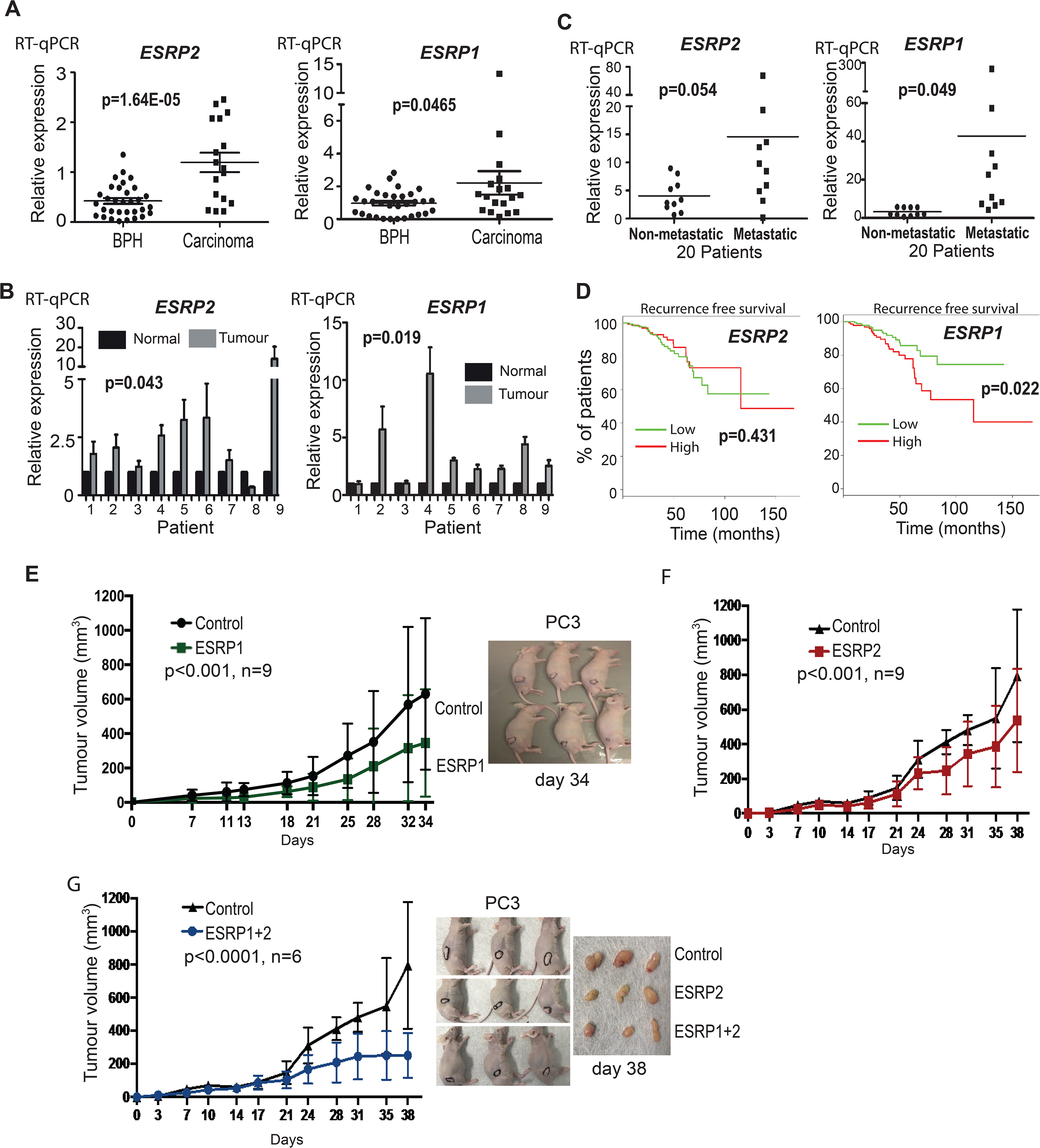
ESRP2 and its paralog ESRP1 are highly expressed in primary prostate tumours. **(A)** Real-time PCR analysis of *ESRP1* and *ESRP2* mRNA from patients with benign prostate hyperplasia (BPH) and 17 malignant samples from transurothelial resection of the prostate (TURP) samples. **(B)** Real-time PCR analysis of *ESRP1* and *ESRP2* mRNA from normal and matched prostate cancer tissue from 9 patients obtained from radical prostatectomy. **(C)** Analysis of *ESRP 1* and *ESRP2* mRNA levels in samples from the Walker et al (51) cohort. Statistical analysis in parts (A)-(C) were performed using t tests. **(D)** Interrogation of the TCGA PRAD (PRostate ADenocarcinoma) cohort using KM-express (52). *ESRP1* expression levels linked to a reduced time to PSA biochemical recurrence (bifurcate gene expression at average, log-rank test p=0.022). Over-expression of **(E)** ESRP1, **(F)** ESRP2, or **(G)** both ESRP1 and ESRP2 in PC3 cells significantly slowed the growth of prostate cancer xenografts *in vivo*. Data were analysed by Two-way ANOVA, and the p value is for the overall difference between two groups.

Each of the above data showed that *ESRP1* and *ESRP2* expression levels are relatively high in primary prostate cancer compared to normal prostate tissue. High *ESRP2* expression was not prognostic of disease progression in the TCGA (PRostate ADenocarcinoma) PRAD cohort using KM-express (52), but high expression of *ESRP1* associated with a significantly reduced time to first biochemical recurrence (p=0.022) (Figure 2D). Previous data have reported up-regulated ESRP1 and ESRP2 proteins in squamous cell carcinoma tumours but their down-regulation at invasive fronts (53). We tested these same antibodies against ESRP1 and ESRP2 proteins on prostate cancer FFPE tissue and cell blocks, but they did not pass our stringent quality control tests (Figure 1-Figure supplement 1C). While this manuscript was in preparation, another group used an alternative ESRP1 antibody to show upregulation of ESRP1 in 12,000 prostate cancer tissue microarray tumours (54).

We next investigated the effects of ESRP1/2 expression on the biology of prostate cancer cells *in vivo*. Because of their normal endogenous expression profiles (Figures 1C and 1D), we selected PC3 and DU145 cells to study the effects of ESRP1/ESRP2 protein up-regulation on prostate cancer cells. Ectopic expression of ESRP1 and ESRP2 protein expression in AR negative PC3 and DU145 cell line models reduced prostate cancer cell growth *in vitro* (Figure 2-Figure supplement 1). Over-expression of both ESRP1 and ESRP2 (either alone or together) in PC3 cells also significantly slowed growth of prostate cancer xenografts *in vivo* (Figures 2E-G). Taken together, the above data show that ectopic expression of ESRP1 and ESRP2 proteins slow the growth of PC3 and DU145 prostate cancer cell lines and are strongly suggestive that high levels of ESRP2 protein inhibit growth of prostate cancer cells.

### Identification of endogenous ESRP1/ESRP2-regulated targets in prostate cancer cells

To enable us to test whether androgens may control splicing indirectly via transcriptional regulation of *ESRP2*, we next set out to identify a panel of endogenous ESRP2-responsive exons within prostate cancer cells. We first used siRNAs to jointly deplete both ESRP1 and ESRP2 proteins from LNCaP cells (since ESRP1 and ESRP2 can regulate overlapping targets); and in parallel treated LNCaP cells with a control siRNA. We then used RNAseq to monitor the effects of these treatments on the LNCaP transcriptome. Bioinformatic (55) analysis of these RNAseq data (GSE129540) predicted 446 ESRP1/ESRP2 regulated alternative splicing events across 319 genes (ΔPSI>10%, p<0.05) (Figure 3-source data 1). We experimentally validated splicing switches for 44 predicted ESRP1/ESRP2-controlled exons by RT-PCR analysis, after LNCaP cells were treated with either of two independent sets of siRNA directed against ESRP1 and ESRP2 or control siRNAs (Figure 3 and Figure 3 source data 2). We also detected similar splicing switches for 37/44 of these skipped exons after jointly depleting ESRP1 and 2 from the AR-positive CWR22 RV1 prostate cancer cell line; and 28/44 of these splicing switches were observed after jointly depleting ESRP1 and 2 from the AR positive PNT2 cells that model the normal prostate epithelium (Figure 3 and Figure 3-source data 2).

**Figure 3.**
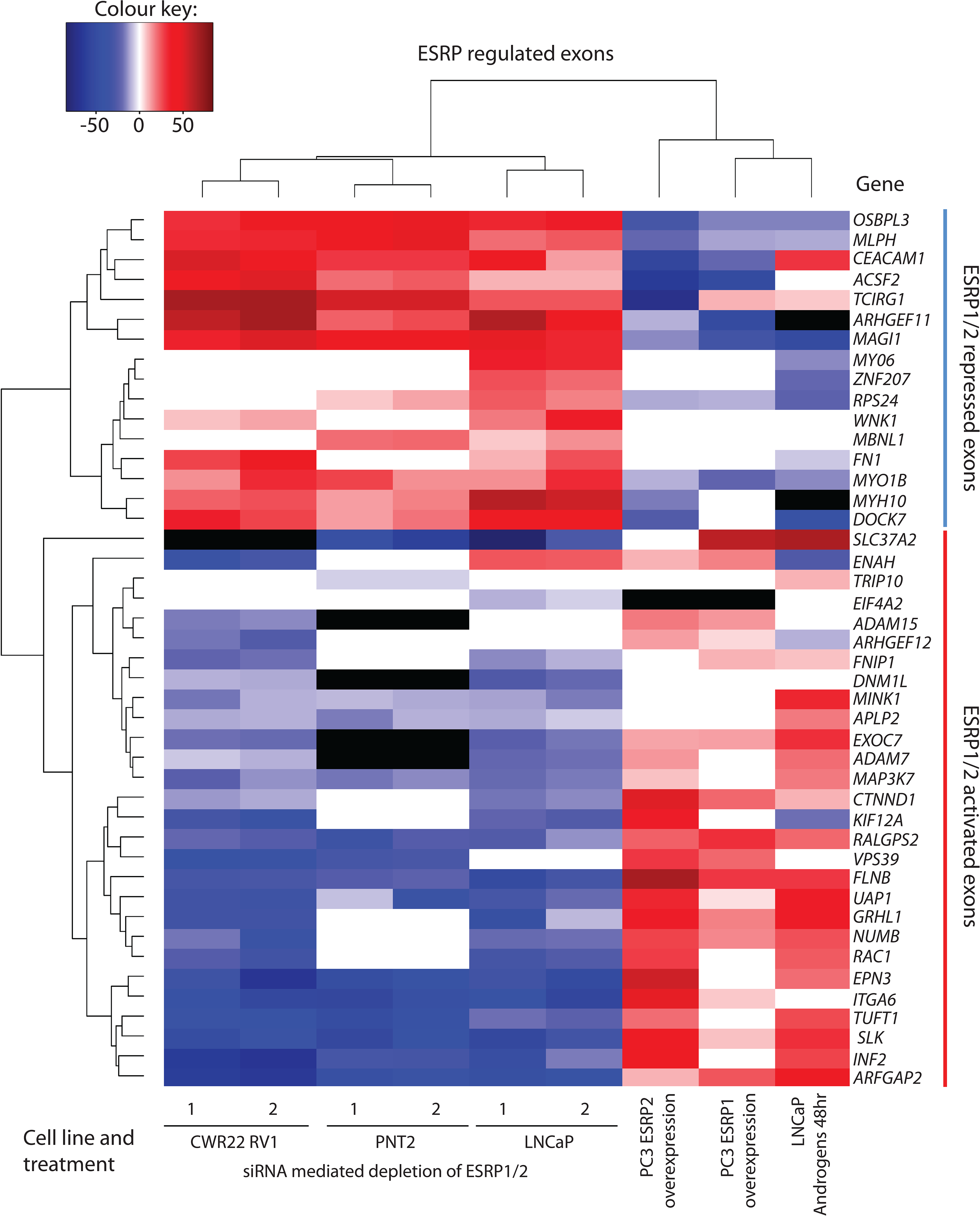
Identification of endogenous ESRP1/ESRP2 regulated target exons in prostate cancer. Heat map showing mean PSI levels for a panel of ESRP-regulated exons in prostate cancer cell lines (CWR22RV1, PNT2, LNCaP and PC3). Mean PSIs were calculated for ESRP-regulated isoforms between cells treated with siRNAs specific to ESRP1 and ESRP2, or control siRNAs (CWR22RV1, PNT2, LNCaP), between PC3 cells with and without ectopic expression of ESRP1 or ESRP2, and between LNCaP cells grown in steroid deplete versus androgen stimulated conditions (10nM R1881 for 48 hours). Biological triplicate samples were used for CWR22RV1, PNT2 and LNCaP cells, and technical replicate samples were used for RNAs prepared from PC3 cells. PSI levels were measured using RT-PCR analysis averaged from 3 replicates (mean data given in Figure 3-source data 2), and clustered in the heat map according to splicing patterns across the different conditions. The heatmap was generated using heatmap 2 function using R’s ‘gplots’ package. The black shading in the heatmap denotes non-detection of the mRNA isoform after RT-PCR, and white denotes no change detected.

Given this set of endogenous target exons, we carried out further analyses to next identify target exons that respond to increasing levels of either ESRP2 or ESRP1 expression in PC3 cells (which normally express low levels of endogenous ESRP1/ESRP2) (Figure 1C). Ectopic expression of either ESRP1 or ESRP2 in PC3 cells induced splicing switches for 35/44 exons analysed. Importantly, the splicing switches induced by ectopic expression of either ESRP2 or ESRP1 were reciprocal to the splicing switches detected after siRNA depletion of ESRP1/ESRP2 (Figure 3). Experimentally validated ESRP-regulated exons fell into two groups: splicing of one group was repressed by ectopic expression of ESRP1 or ESRP2 in PC3 cells, and reciprocally activated by endogenous ESRP1/ESRP2 depletion in LNCaP cells (these exons are in the top of the heatmap in Figure 3, from *OSBL3* to *DOCK7*); and a second group whose splicing was activated by ectopic expression of ESRP1 or ESRP2, and reciprocally repressed by ESRP1/ESRP2 depletion (from *SLC37A2* to *ARFGAP2* in Figure 3).

### An androgen steroid hormone-ESRP2 axis controls alternative splicing in AR-positive prostate cancer cells

The above data thus identified a robust panel of alternative exons within prostate cancer cells that responded to ESRP1/ESRP2 expression levels. We next tested if this panel of ESRP2-regulated exons are also regulated by ambient androgen concentrations. LNCaP cells were harvested after growth in steroid deplete media and after 48 hours of androgen stimulation (this timing was designed to enable full levels of androgen-mediated ESRP2 protein induction, Figure 1G). Our prediction was that androgen stimulation of LNCaP cells would activate ESRP2 expression to regulate our panel of endogenous test exons. If this was the case, splicing switches in response to androgen stimulation should occur in a reciprocal direction to splicing changes induced by ESRP1/ESRP2 protein depletion in LNCaP cells. Consistent with these expectations, more than 70% (32/42) exons in our test panel demonstrated androgen regulated splicing (Figure 3-source data 2). Importantly, plotting the percent spliced-in (PSI) for each exon after 48 hours androgen stimulation (Y axis) versus the PSI after ESRP1/ESRP2 depletion (X axis) showed a significant negative correlation (slope= −0.66, R^2^ = 0.64, p<0.0001) (Figure 4A).

**Figure 4.**
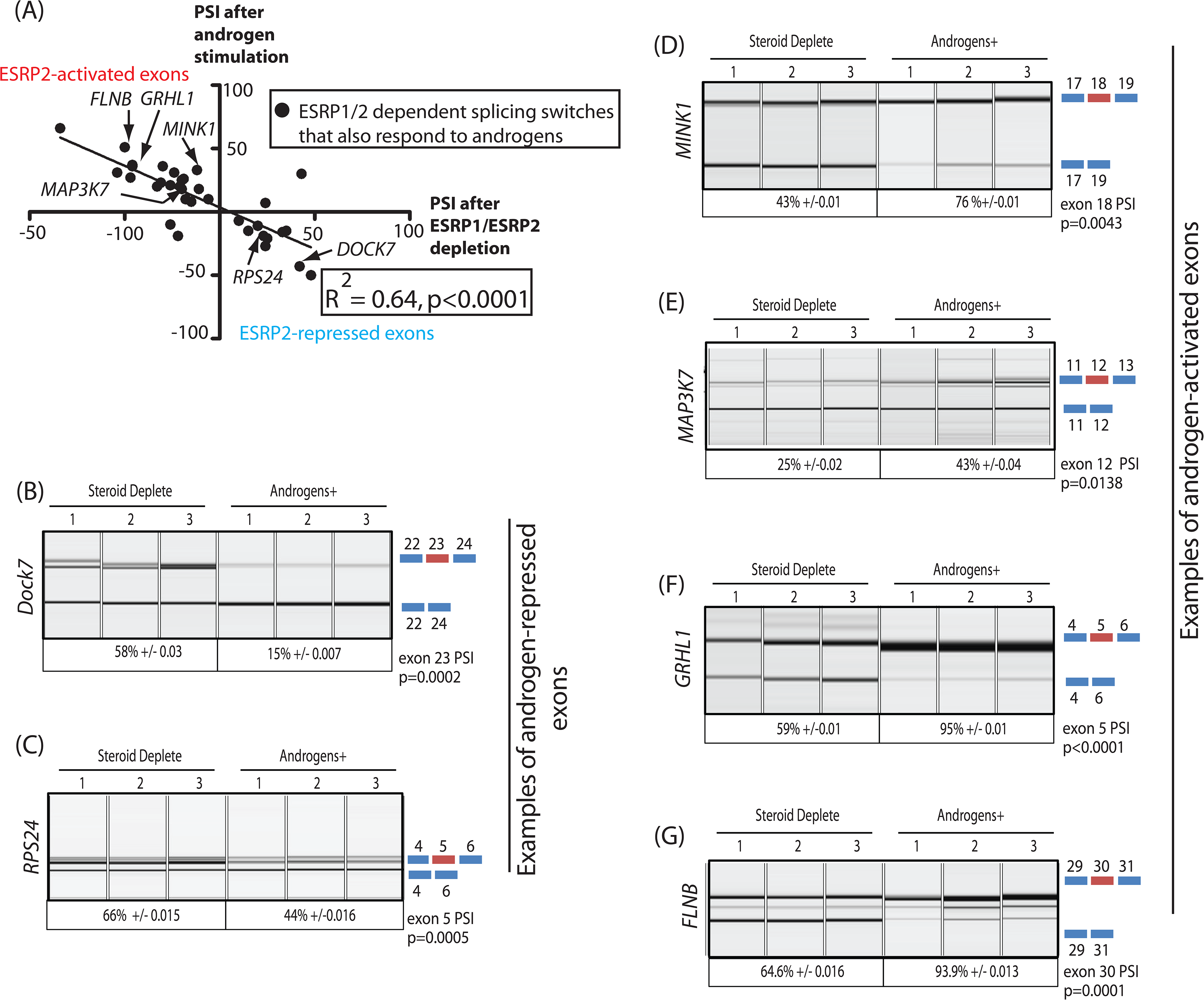
An androgen steroid hormone-ESRP2 axis controls alternative splicing in prostate cancer cells. **(A)** ESRP2-regulated exons are also frequently controlled by androgens in prostate cancer cells. 31/48 of the ESRP target exons (identified by RNAseq analysis of LNCaP cells depleted of ESRP1 and ESRP2) were regulated in the opposite direction in LNCaP cells treated by androgens (10nM R1881) for 48 hours (which would induce ESRP2 protein expression). Plotting the splicing responses to androgen stimulation with those after ESRP1/ESRP2 depletion revealed a negative correlation (slope= −0.66+/-0.09, Rsquare=0.64, p<0.0001, calculated using Graphpad). Individual values for this graph are given in Figure 3 – source data 2, and are averages from 3 biological replicates. **(B-C)** Capillary gel electrophoretograms showing splicing patterns of 3 biological replicates grown in steroid deplete media, or steroid deplete media supplemented with R1881, for alternative exons in the **(B)** *DOCK7* and **(C)** *RPS24* genes that were repressed by 48 hours androgen treatment. **(D-G)** Capillary gel electrophoretograms showing splicing patterns in 3 biological replicates grown in steroid deplete media, or steroid deplete media supplemented with R1881, for alternative exons in the **(D)** *MINK1*, **(E)** *MAP3K1*, **(F)** *GRHL1* and (**G**) *FLNB* genes that were activated by 48 hours of androgen treatment. For parts (B)-(G) the p values were calculated using unpaired t tests.

These results experimentally support an androgen-ESRP2 axis controlling splicing patterns in prostate cancer cells. Amongst the genes containing ESRP2-repressed exons that were also skipped in response to androgen stimulation were *DOCK7* (exon 23), which encodes a guanine nucleotide exchange factor involved in cell migration (Figure 4B) (56); and *RPS24* (exon 5), a gene that is highly expressed in prostate cancer (Figure 4C) (57). Amongst the genes containing ESRP-activated exons that were also activated by androgen exposure were *MINK1* (exon 18) which encodes a pro-migratory serine/threonine kinase (Figure 4D); *MAP3K7* (exon 12) which encodes a serine/threonine kinase that regulates signalling and apoptosis, activates NFKappaB, and is lost in aggressive prostate cancer (58, 59) (Figure 4E); *GRLH1* (exon 5) that encodes a transcription factor involved in epithelial cell functions (60) (Figure 4F); and *FLNB* (exon 30), alternative splicing of which has been identified as a key switch contributing to breast cancer metastasis (61, 62) (Figure 4G).

### The AR-ESRP2 axis controls splicing of mRNA isoforms that are important for prostate cancer disease progression

Information about most of our panel (38/44) of ESRP-regulated exons was also found within the TCGA PRAD cohort (containing 497 prostate tumour samples and 52 samples from normal prostate tissue). Analysis of the PRAD cohort revealed that 18/38 ESRP-regulated exons have different patterns of splicing inclusion between tumour and normal tissue (Figure 5A and Figure 3 source data 2). These differentially spliced exons include the AR-ESRP2-controlled alternative exons in the *DOCK7* and *RPS24* genes (both of which were excluded in prostate tumours compared to normal prostate tissue); and the alternative exons in the *MINK1* and *MAP3K7* genes (each of which had increased levels of splicing inclusion in prostate tumours compared to normal tissue). Further RT-PCR analysis of an independent cohort confirmed more frequent skipping of *DOCK7* (exon 23) and *RPS24* (exon 5) in prostate tumour tissue compared to normal prostate (Figure 5B and 5C).

**Figure 5.**
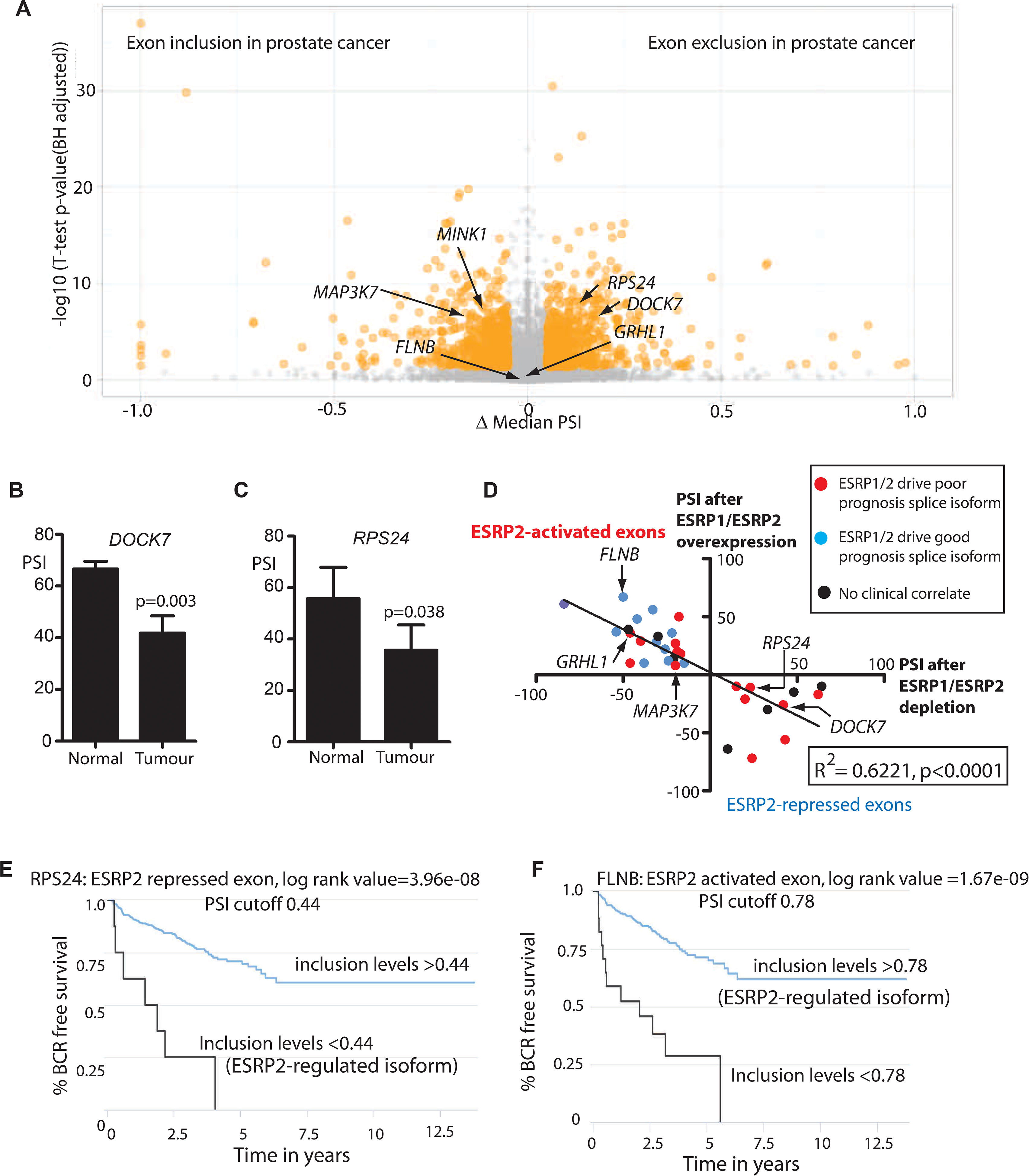
Alternative splicing patterns controlled by the androgen steroid hormone-ESRP2 splicing axis are clinically relevant for disease progression. **(A)** Volcano plot showing PSIchomics (76) alternative splicing analysis of RNAseq data performed between normal prostate tissue and prostate tumour tissue from the TCGA PRAD cohort (consisting of 497 prostate tumour samples and 52 normal tissue). The t-test p-value (Benjamini-Hochberg adjusted for multiple testing) was used as metric of statistical significance. **(B)** Percentage splicing inclusion, quantified by RT-PCR, of *DOCK7* exon 23 within samples of prostate tumour and adjacent normal tissue (statistical significance calculated using t test). **(C)** Percentage splicing inclusion, quantified by RT-PCR, of *RPS24* exon 2 within 9 matched samples of prostate tumour and adjacent normal tissue (statistical significance calculated using t test). **(D)** Graphical representation of levels of average PSI levels in response to ectopic ESRP2 expression in PC3 cells (Y axis) versus after ESRP1/ESRP2 depletion in LNCaP cells. Individual PSI values to make this graph are averaged from 3 biological replicates, and are given in Figure 3 – source data 2. Note that the PSI values for ESRP over-expression refer to ESRP2 over-expression, with the exception of *FNIP1* and *SLC37A2* that are for ESRP1 over-expression (see Figure 3 – source data 2). Linear regression analysis of this data was analysed using Graphpad. Individual splice forms were correlated with clinical data for time to PSA biochemical recurrence within the PRAD cohort (76). Points on this graph corresponding to individual ESRP-regulated splice isoforms are coloured differently according to whether they correlated with an increased time to PSA biochemical recurrence (blue dots), a decreased time to biochemical recurrence (red dots) or had no significant correlation (black dots) is shown. **(E-F)** Kaplan-Meier plots showing data from TCGA PRAD cohort of percentage of tumours that are free of biochemical recurrence versus time in years, associated with expressing the alternative splice isoforms of (E) *RPS24* exon 5 (PSI cut off 0.44), and (F) *FLNB* exon 30 (PSI cut off 0.78) (76).

To visualise the amplitude of splicing switches in these exons in response to ESRP2, we plotted PSIs measured *in vitro* after ectopic expression of ESRP1/ESRP2, versus PSI values after siRNA mediated depletion of ESRP1/ESRP2 (Figure 5D, using data from Figure 3 and Figure 3 - source data 2, slope= −0.74, R^2^ = 0.6221, p<0.0001). We then monitored the data in TCGA PRAD cohort for time taken to first biochemical tumour recurrence associated with splicing inclusion for each of these exons. This revealed 3 groups of ESRP-regulated exons, that either correlated with an increased or decreased time to recurrence, or alternatively showed no correlation (individual plots are shown in Figure 5 –figure supplement 1). Splicing inclusion ESRP1/ESRP2-controlled exons associated with an increased time to biochemical recurrence are shown in blue in Figure 5D. This group included *FLNB* exon 30, which also had the highest amplitude splicing inclusion level observed in response to ESRP2 expression in PC3 cells. *FLNB* exon 30 was also strongly skipped after siRNA depletion of ESRP1/ESRP2 in LNCaP cells (Figures 3 and 5D) and strongly activated in response to androgen stimulation (Figure 4G). In the PRAD dataset, levels of *FLNB* exon 30 splicing inclusion above 0.78 correlated with an increased time to biochemical recurrence, so a more favourable clinical outcome in prostate cancer (Figure 5F). Splicing inclusion of the second set of ESRP1/ESRP2-regulated exons that correlated with decreased time to biochemical recurrence are shown in red in Figure 5D. These included exons both activated (*GRHL1* exon 5 and *MAP3K7* exon 12) and repressed by the AR-ESRP2 axis (*DOCK7* exon 23, and *RPS24* exon 5D and 5E). Splicing inclusion of the third and smallest set of exons that did not correlate with time to biochemical recurrence are identified with black dots in Figure 5D.

### Splicing of a key exon in the *FLNB* gene is switched by a drug that antagonises AR activity

The above data identified a subset of ESRP2-regulated splicing switches that associated with biochemical recurrence of prostate cancer after treatment. Since ESRP2 expression was repressed in patient prostate cancer tissue by ADT, we next investigated whether AR inactivation may inadvertently modulate splice isoforms in genes important for cancer progression. We focussed this analysis on skipping of *FLNB* exon 30, which has recently reported to be a key driver of EMT in breast cancer development (62). Androgen induction of *ESRP2* mRNA expression was blocked by the androgen antagonist bicalutamide (Casodex^®^) (Figure 6A). Consistent with our prediction, treatment of LNCaP cells with Casodex^®^ also reduced splicing inclusion levels of *FLNB* gene exon 30 by almost 20% (Figure 6B). ESRP2 protein expression was also reduced by siRNA depletion of the AR (Figure 6C). Furthermore, siRNA-mediated depletion of AR also significantly reduced levels of *FLNB* splicing inclusion from 84% to 69% (Figure 6D). Both these data support a scenario where splicing inclusion of *FLNB* gene exon 30 is modulated in response to ADT as well as androgen stimulation.

**Figure 6.**
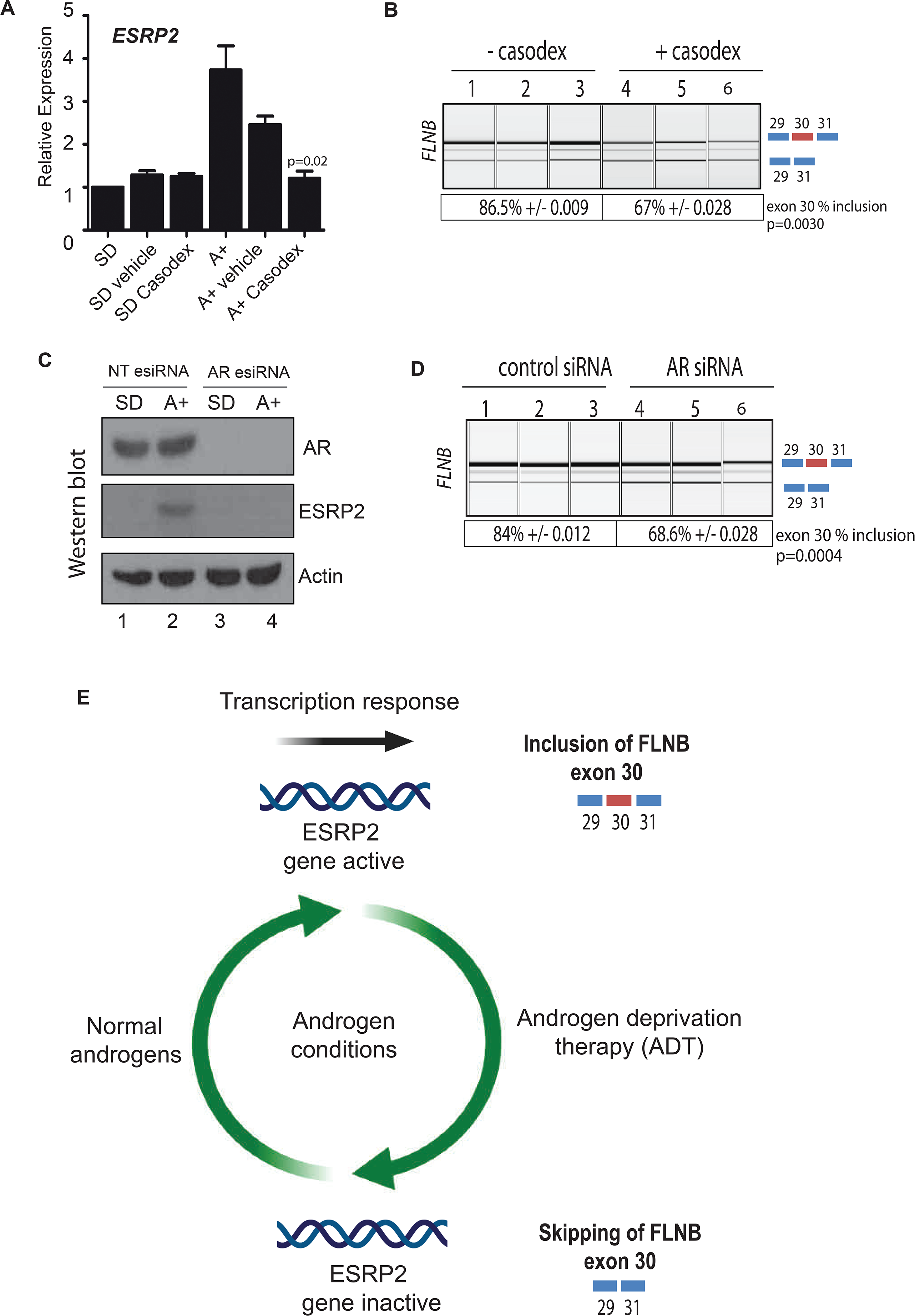
Inhibition of AR function switches ESRP2-dependent splicing patterns. **(A)** *ESRP2* mRNA expression in cells grown in steroid deplete (SD) conditions, and after addition of androgens (A+)(quantified by real-time PCR from three biological replicates). Androgen-mediated activation was inhibited in the presence of 10 μM of the anti-androgen bicalutamide (Casodex^®^). Cells were cultured for 24 hours. Statistical significance was calculated using a t test between the A + vehicle and the A + casodex samples. **(B)** Capillary gel electrophoretogram showing RT-PCR analysis measuring splicing inclusion levels for *FLNB* exon 30 +/-24 hours Casodex^®^ treatment (three biological samples shown, statistical significance calculated using a t test). **(C)** Western blot showing levels of the AR, ESRP2 and actin in samples of LNCaP cells following growth in steroid deplete media (SD), or plus androgens for 48 hours (A+). Cells were transfected with esiRNAs designed to deplete the AR, or a control siRNA (NTesiRNA). Statistical significance was calculated using a t test. **(D)** Capillary gel electrophoretogram showing RT-PCR analysis of 3 biological replicate RNA samples, measuring splicing inclusion levels for *FLNB* exon 30 after siRNA depletion of the AR or treatment with a control siRNA. **(E)** Model describing how exposure to androgens regulates splicing patterns in prostate cancer cells. Androgen exposure leads to transcription of the gene encoding the splicing regulator protein to promote epithelial splicing patterns, including splicing inclusion of *FLNB* exon 30. ADT leads to transcriptional repression of ESRP2, leading to a dampening of epithelial splicing patterns, including inclusion of *FLNB* exon 30. Image created using BioRender.

## Discussion

In this study we report a novel molecular mechanism to explain how androgen steroid hormones control splicing patterns in prostate cancer cells, and that unifies the functions of the AR both as a transcription factor and being able to control splicing. In this model, the AR controls expression of the master splicing regulator protein ESRP2, which then regulates the splicing patterns of key genes important for prostate cancer biology (Figure 6E). Amongst the key data supporting this proposed model, we find that *ESRP2* is a direct and early target for transcriptional activation by the AR in prostate cancer cells. Furthermore endogenous splice isoform patterns controlled by ESRP1 and ESRP2 target also respond to androgen stimulation, siRNA-mediated depletion of the AR and/or the AR inhibitor bicalutamide (Casodex^®^). While intuitively straightforward, this model is conceptually different from the mechanisms through which estrogen and progesterone have been shown to regulate splicing (via recruitment of splicing regulators as transcriptional cofactors, and by modulation of transcription speeds and chromatin structure).

Androgens are already known to substantially modify the prostate cancer transcriptome at the transcriptional level, with important implications for cell behaviour and cancer progression (29). The data presented here imply that androgens also have an important role in controlling splicing patterns, particularly those that relate to epithelial functions. Previous studies identified just a small number of alternative exons that are controlled by androgens in prostate cancer cells, none of which overlapped with the current study (9, 11). We suggest that an important reason for this discrepancy is because previously splicing patterns were monitored after 24 hours of androgen exposure. Since we now show that splicing regulation by androgens operates indirectly through transcriptional control of ESRP2, 24 hours androgen exposure would not be sufficient to upregulate ESRP2 levels. For the panel of exons we have investigated in the current study, we analysed androgen-dependent splicing switches after 48 hours, to allow sufficient time for ESRP2 induction at the protein level and re-equilibration of downstream splice isoform ratios.

The expression of ESRPs appears to be plastic during cancer progression (36, 53, 63) and ESRPs have previously been shown to have a dual role in carcinogenesis with both gain and loss associated with poor patient prognosis (36). ESRP1 has recently been shown to be amplified in an aggressive subgroup of early onset prostate cancer (54). ESRP1 expression is linked to poor survival and metastasis in lung cancer (64), and both ESRP1 and ESRP2 are upregulated in oral squamous cell carcinoma relative to normal epithelium (53). However, analyses of clinical datasets imply that the expression levels of ESRP2 in patients may not be prognostic in themselves for prostate cancer progression. Instead, because ESRP2 is a critical component of epithelial-specific splicing programmes, we suggest that down-regulation of ESRP2 levels in response to ADT could be of importance in prostate cancer patients, since this will dampen epithelial splicing patterns, helping to prime prostate cancer cells for future mesenchymal development and possible metastasis.

*FLNB* exon 30 was amongst the highest amplitude splicing switches detected in response to ESRP2 expression in PC3 cells and androgen stimulation in LNCaP cells. Supporting the possibility that clinically important splice isoforms may switch in response to ADT, *FLNB* exon 30 skipping was also significantly increased by both bicalutamide (Casodex^®^) treatment and siRNA depletion of the AR. *FLNB* encodes an actin binding protein which is linked to cancer cell motility and invasion (65, 66). Importantly, this same splicing switch in *FLNB* exon 30 is sufficient to initiate metastatic progression in breast cancer (62). The clinical prognosis of metastatic prostate cancer is poor (4). This makes the mechanisms that control metastasis of prostate cancer cells, and any links with ADT of prime importance. In prostate cancer EMT has been linked to a common mechanism underlying therapeutic resistance and is associated with poor prognosis (16). Sun et al. showed that although ADT can effectively control prostate tumour size initially, it simultaneously promotes EMT, an unintended consequence that could ultimately lead to CRPCa (21). Such direct links between ADT and EMT uncover an important yet overlooked consequence of the standard care treatment for prostate cancer (67). Although the causes of EMT in prostate cancer progression to CRPCa are likely to be complex, the down-regulation of ESRP proteins has been shown to be essential for EMT progression (68). Thus loss of ESRP expression may provide a molecular explanation why AR positive prostate cancer cells show increased susceptibility to EMT in response to ADT, and so is relevant to consider with regard to therapy. Our findings have important implications for second line treatment strategies in a clinical setting, and suggest an alternative approach may be to inhibit EMT in combination with ADT to prevent disease progression.

## Materials and Methods

### Cell Culture

Cell culture and androgen treatment of cells was as described previously (11, 69-73). All cells were grown at 37°C in 5% CO2. LNCaP cells (CRL-1740, ATCC) were maintained in RPMI-1640 with L-Glutamine (PAA Laboratories, R15-802) supplemented with 10% Fetal Bovine Serum (FBS) (PAA Laboratories, A15-101). For androgen treatment of LNCaP cells, medium was supplemented with 10% dextran charcoal stripped FBS (PAA Laboratories, A15-119) to produce a steroid-deplete medium. Following culture for 72 hours, 10nM synthetic androgen analogue methyltrienolone (R1881) (Perkin–Elmer, NLP005005MG) was added (Androgen +) or absent (Steroid deplete) for the times indicated.

### Antibodies

The following antibodies were used for western blotting: Anti-ESRP2 rabbit antibody (Genetex, GTX123665), anti-rabbit ESRP1 (Sigma, HPA023719), anti-AR mouse antibody (BD Bioscience, 554226), anti-actin rabbit antibody (Sigma, A2668), anti-FLAG mouse monoclonal antibody (Sigma, F3165), normal rabbit IgG (711-035-152 Jackson labs) and normal mouse IgG (715-036-150 Jackson labs). For immunohistochemistry the following ESRP antibodies were tested: anti-rabbit ESRP1 (Sigma, HPA023719) and anti-rabbit ESRP2 (Abcam ab113486) but were found not to be specific for FFPE cell pellets.

### RT-qPCR

Cells were harvested and total RNA extracted using TRI-reagent (Invitrogen, 15596-026), according to the manufacturer’s instructions. RNA was treated with DNase 1 (Ambion) and cDNA was generated by reverse transcription of 500ng of total RNA using the Superscript VILO cDNA synthesis kit (Invitrogen, 11754-050). Quantitative PCR (qPCR) was performed in triplicate on cDNA using SYBR® Green PCR Master Mix (Invitrogen, 4309155) using the QuantStudio™ 7 Flex Real-Time PCR System (Life Technologies). ESRP1 was detected using (ESRP1 for AGCACTACAGAGGCACAAACA; ESRP1 Rev TGGAGAGAAACTGGGCTACC). ESRP2 was detected using the primer combination (ESRP2 For CCT GAA CTA CAC AGC CTA CTA CCC; ESRP2 Rev TCC TGA CTG GGA CAA CAC TG). Samples were normalised using the average of three reference genes: GAPDH (GAPDH For AAC AGC GAC ACC CAT CCT C; GAPDH Rev TAGCACAGCCTGGATAGCAAC); β –tubulin (TUBB For CTTCGGCCAGATCTTCAGAC; TUBB Rev AGAGAGTGGGTCAGCTGGAA); and actin (ACTIN For CATCGAGCACGGCATCGTCA; ACTIN Rev TAGCACAGCCTGGATAGCAAC).

### siRNA

siRNA mediated protein depletion of ESRP1/2 was carried out using Lipofectamine RNAiMAX Transfection Reagent (Thermo Fisher, 13778075) as per the manufacturer’s instructions and for the times indicated. The siRNA sequences used were ESRP1 siRNA1 (hs.Ri.ESRP1.13.1); ESRP1 siRNA2 (hs.Ri.ESRP1.13.2); ESRP2 siRNA 1 (hs.Ri.ESRP2.13.1); ESRP2 siRNA 2 (hs.Ri.ESRP2.13.2); and a negative control siRNA (IDT (51-01-14-04)). AR esiRNA was as described previously (29).

### Immunohistochemistry

Freshly cut tissue sections were analysed for immunoexpression using Ventana Discovery Ultra autostainer (Ventana Medical Systems, Tucson, Arizona). In brief, tissue sections were incubated in Cell conditioning solution 1 (CC1, Ventana) at 95°C to retrieve antigenicity, followed by incubation with respective primary antibodies described above. Bound primary antibodies were visualized using UltraMap DAB anti-Rb Detection Kit.

### AR-ChIP

LNCaP cells were stimulated with 10 nM R1881 overnight. The ChIP assay was performed using the one step ChIP kit (Abcam ab117138) as per manufacturer’s instruction. Briefly, cells were fixed and crosslinked in 1% formaldehyde for 10 minutes at 37 °C and incubated with protease inhibitors. Chromatin was isolated from cell lysates and enzymatically fragmented using an EZ-Zyme Chromatin Prep Kit (Merck 17 375). 10 ug of anti - AR antibody (Abcam ab74272) or IgG control antibody was used to precipitate DNA crosslinked with the androgen receptor. Enriched DNA was then probed by qPCR using primers targeting the ESRP2 regulatory region to assess AR binding intensity. Primer sequences used to detect PSA were (PSA ChIP for GCC TGG ATC TGA GAG AGA TAT CAT C; PSA Chip rev ACA CCT TTT TTT TTC TGG ATT GTT G). Primers used to detect AR binding near to ESRP2 were (ESRP2 Chip for TCCCGAGTAGCTGGGACTAC; ESRP2 Chip rev CAGTGGCTTACACCTGGGAG).

### Creation of PC3 stable cell lines

The ESRP1 plasmid (PIBX-C-FF-B-ESRP1) was a gift from Prof Russ Carstens (University of Philadelphia. USA) and the ESRP2 plasmid (pBIGi hESRP2-FLAG) from Dr Keith Brown (University of Bristol, UK). PC3 cells were transfected using FuGene^®^ HD Transfection Reagent as per manufacturer’s instructions. Stable transfectants with ESRP1 was selected using 10µg/ml Blasticidin and ESRP2 plasmid was selected using 150ug/ml Hygromycin. ESRP2 Plasmid was inducible by 2.5ug/ml doxycycline for 48 hours. PC3 ESRP1 overexpressed cells were transfected with pBIGi hESRP2-FLAG plasmid using the same protocol.

### *In vitro* cell proliferation analysis

For cell growth curves (carried out for *in vitro* analysis of PC3 stable cell lines), PC3 cells were seeded 100,000 cells per well in 12-well plate in 8 plates. Cells were counted every 24 hours after seeding in the plate. All the treatments had 12 repeats. WST assays were carried out over 7 days as per manufacturer’s instructions (Cayman, CAY10008883). For DU145 cells 10,000 cells were seeded per well in a 96 well plate. All data was tested by two-way ANOVA.

### RNAseq analysis

LNCaP cells (passage 19) were treated with either control siRNAs or siRNAs targeting ESRP1 and ESRP2 for 72 hours (samples prepared in triplicate). RNA was extracted 72 hours after siRNA treatment using the Qiagen RNAeasy kit (Cat No. 74104) as per the manufacturer’s instructions. RNAseq was carried out using TruSeq Stranded mRNA Sequencing NextSeq High-Output to obtain 2 × 75 bp reads. Quality control of reads was performed using FastQC. Reads were mapped to the hg38 transcriptome using Salmon. Differential gene expression analysis was performed using DESeq2. Percent spliced-in (PSI) estimates for splicing events were calculated using SUPPA2 (55) based on isoform transcripts per million (TPM) estimates from Salmon (74). Quantification utilised Gencode gene models (release 28). Differential PSI was calculated using DiffSplice using the empirical method (75). Events with a delta PSI?>?10% and FDR?<?0.05 were considered as significant.

### PSIchomics and Bioinformatic analysis of PRAD cohort

Clinical expression patterns of ESRP2-regulated exons were monitored using PSIchomics (76). Differential splicing analysis between primary solid tumour and solid tissue normal samples were subsequently performed to evaluate relative higher inclusion levels in either tumour or normal tissue samples using Δ median and t-test p-value (Benjamini-Hochberg adjusted) values. Survival analysis based on TCGA clinical data derived from prostate cancer patient samples was performed, with time to first PSA biochemical recurrence being the event of interest.

### Tumour xenografts

Stable overexpression of *ESRP1* and stable doxycycline-inducible overexpression of either *ESRP2* alone or *ESRP1* and *2* were obtained using PC3 cells (that have the low endogenous levels of both proteins). One million PC3 overexpressing *ESRP1* or control cells were injected subcutaneously in the flank of male nude mice and tumour volumes were monitored. Two million PC3 cells overexpressing *ESRP2*, overexpressing *ESRP1* and *2*, or control cells were injected subcutaneously in the flank of male nude mice and tumour volumes were monitored. PC3 ESRP2 and PC3 ESRP1/2 cells were cultured in medium supplemented with 2.5ug/ml doxycycline for 48 hours prior to injecting into nude mice to induce ESRP2 expression and mice were administered Doxycycline repeatedly. Tumour diameters were measured by calipers.

### Clinical samples

Our study made use of RNA from 32 benign samples from patients with benign prostatic hyperplasia (BPH) and 17 malignant samples from transurethral resection of the prostate (TURP) samples. Malignant status and Gleason score were obtained for these patients by histological analysis. We also analysed normal and matched PCa tissue from 9 patients obtained by radical prostectomy. The samples were obtained with ethical approval through the Exeter NIHR Clinical Research Facility tissue bank (Ref: STB20). Written informed consent for the use of surgically obtained tissue was provided by all patients. The RNA samples analysed in Figure 2C were previously published (51).

### Statistical Analyses

All statistical analyses were performed using GraphPad Prism 6 (GraphPad Software, Inc.). Statistical analyses were conducted using the GraphPad Prism software (version 5.04/d). PCR quantification of mRNA isoforms was assessed using the unpaired student’s t-test. Data are presented as the mean of three independent samples ± standard error of the mean (SEM). Statistical significance is denoted as * p<0.05, ** p<0.01, *** p<0.001 and **** p<0.0001.

## Acknowledgements

This work was funded by Prostate Cancer UK [PG12-34, S13-020 and RIA16-ST2-011] and the BBSRC [BB/P006612/1]. The work performed at the Vancouver Prostate Centre was funded by the Terry Fox Research Institute (TFRI-NF-PPG). The authors would like to thank Dr Steven Walker, Dr Gemma Logan and Professor Richard Kennedy for kindly providing prostate cancer RNA samples from their 2017 *European Urology* (51) paper for use in Figure 2. The ESRP1 plasmid was a gift from Prof Russ Carstens (University of Philadelphia. USA) and the ESRP2 plasmid from Dr Keith Brown (University of Bristol, UK). Collection of patient RNA samples was supported / funded by the NIHR Exeter Clinical Research Facility, but the opinions given in this paper do not necessarily represent those of the NIHR, the NHS or the Department of Health. The authors thank Dr. Prabhakar Rajan (Barts Cancer Institute, London) for very helpful comments on the manuscript.

---

**Figure 1-Figure Supplement 1.** Confirmation of the specificity of antibodies against ESRP1 and ESRP2. (A) Detection of proteins in PC3 cells by Western blot. (B) Detection of proteins in LNCaP cells by Western blot. (C) Detection of proteins in PC3 cells by immunohistochemistry.

**Figure 1-Figure supplement 1.**
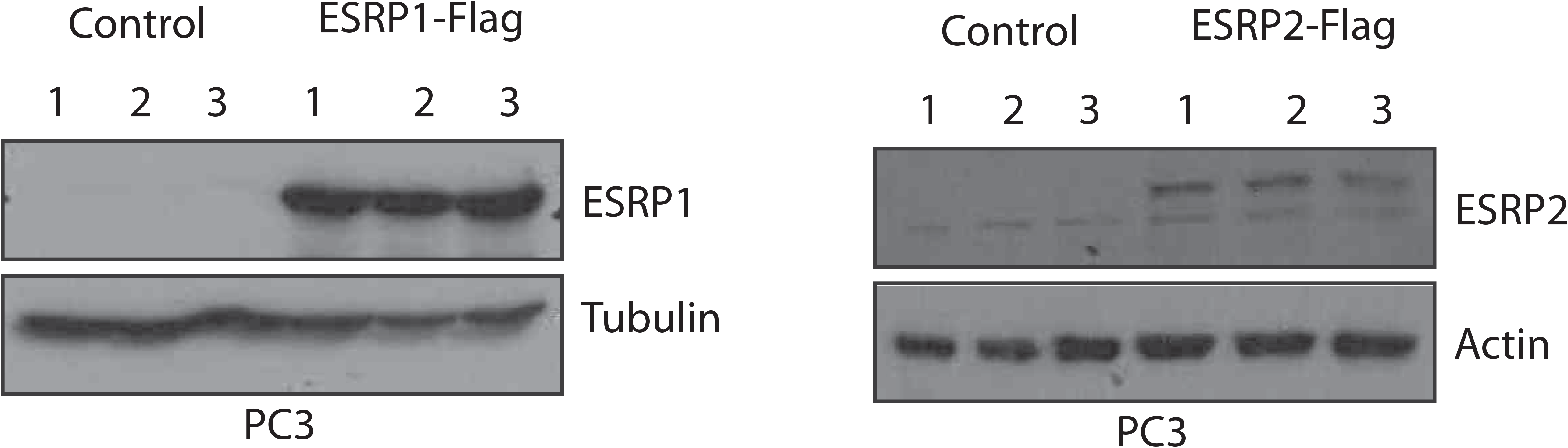
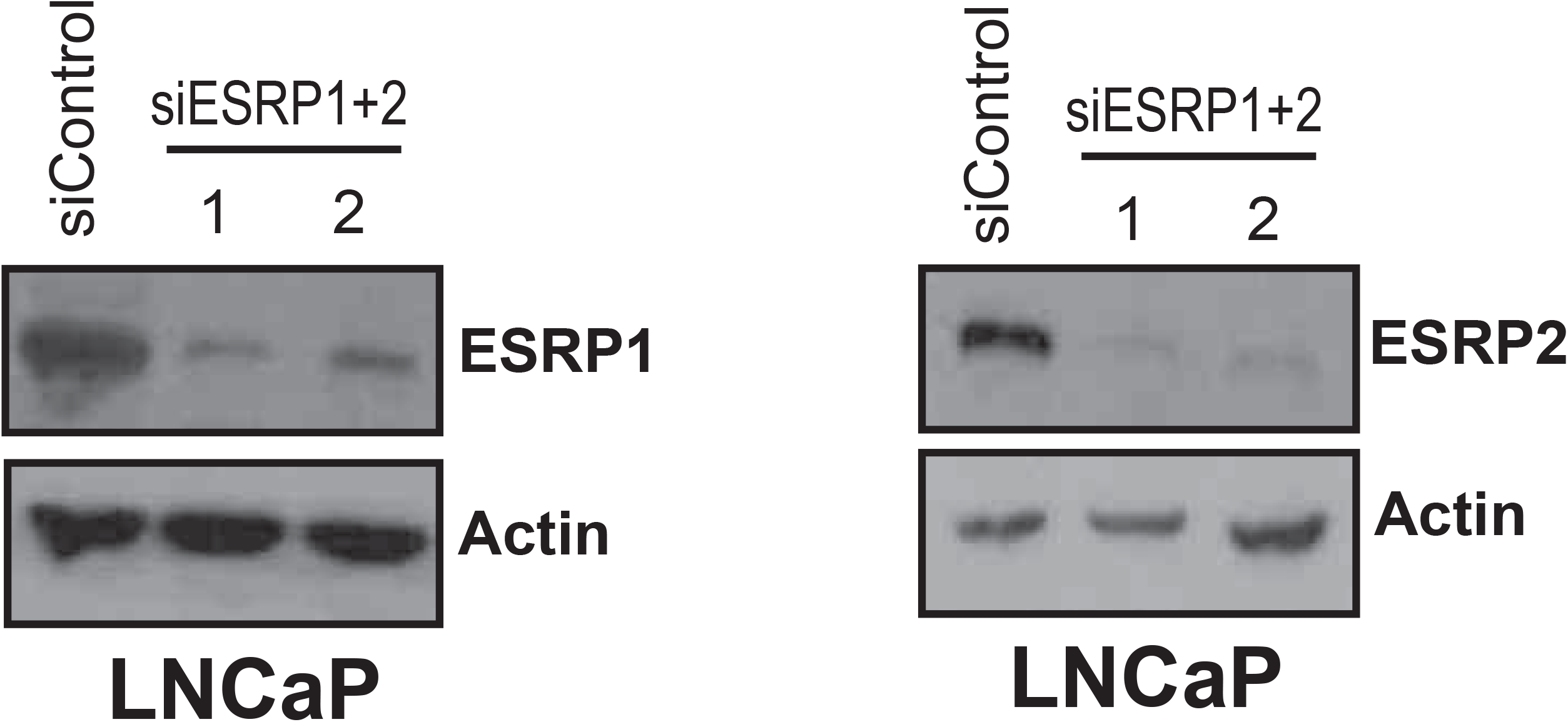
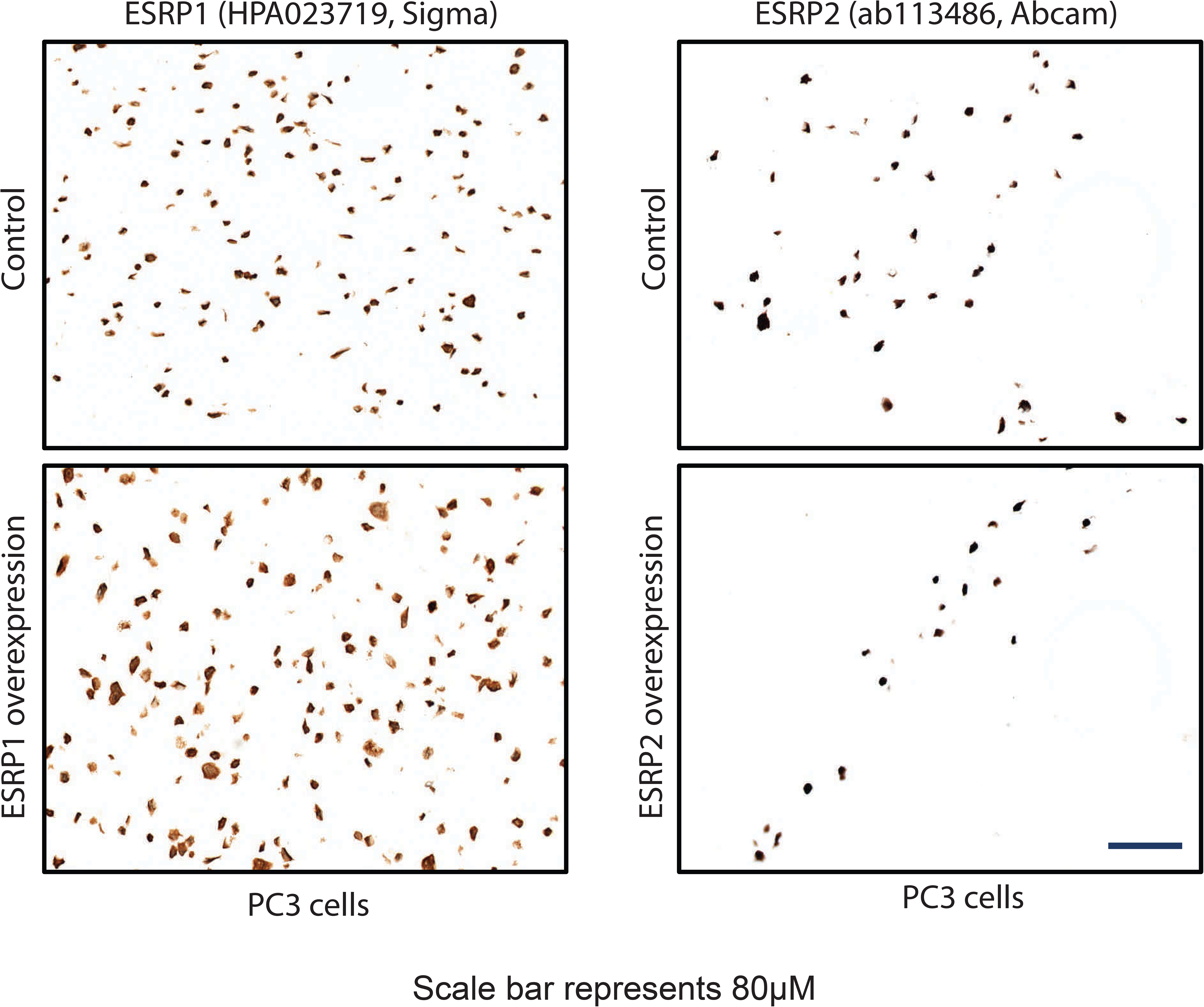
Antibody validation by western blot and IHC A. Detection of overexpressed protein by western blot B. Detection of siRNA mediated protein depletion by western blot C. Detection of overexpressed protein in FFPE cell pellets by IHC

**Figure 2 – Figure Supplement 1.** Ectopic expression of ESRP1 and ESRP2 protein expression in AR negative (A) PC3 and (B) DU145 cell line models reduced prostate cancer cell growth *in vitro*. Data were analysed by Two-way ANOVA, and the p value is for the overall difference between two groups.

**Figure 2-figure supplement 1.**
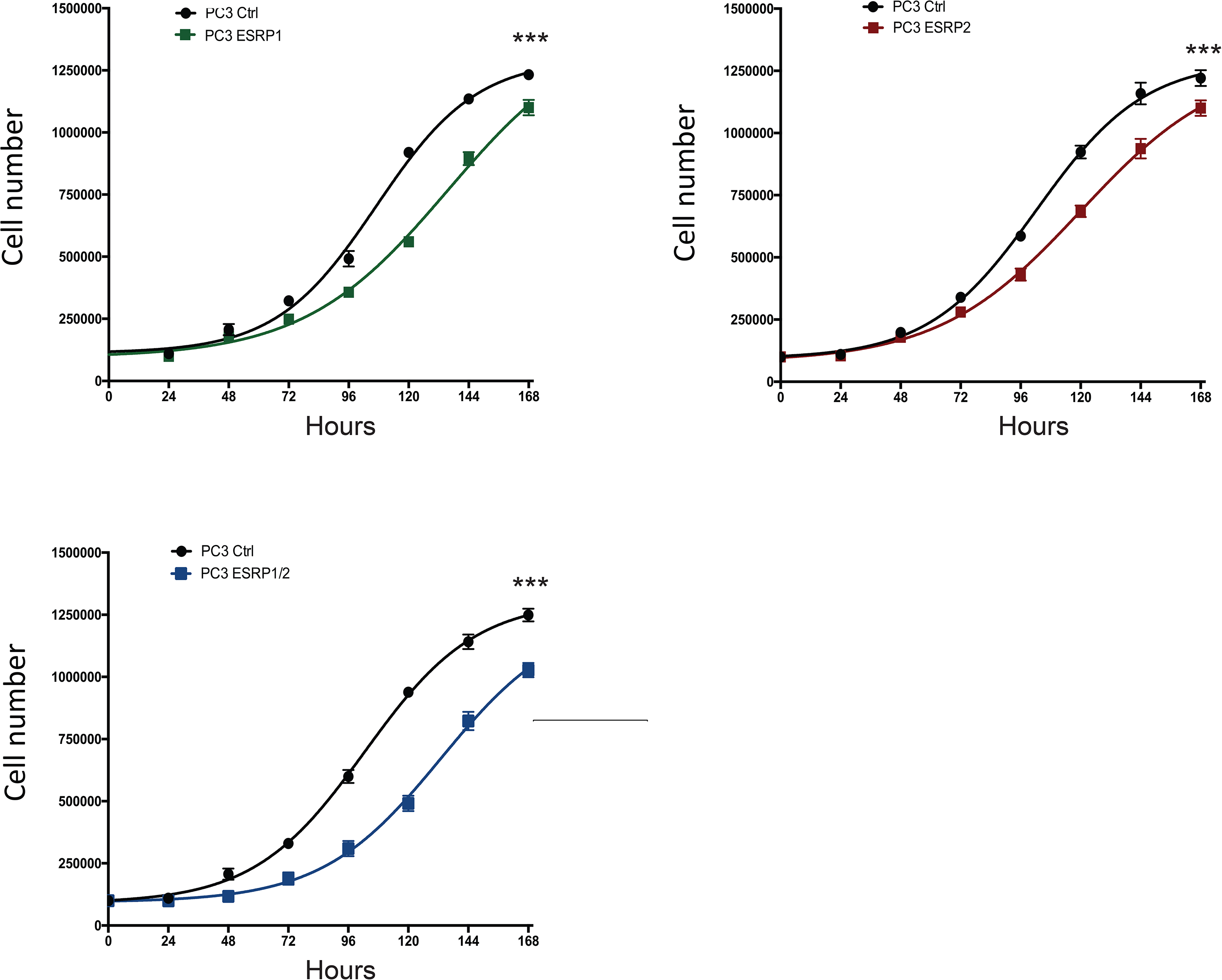
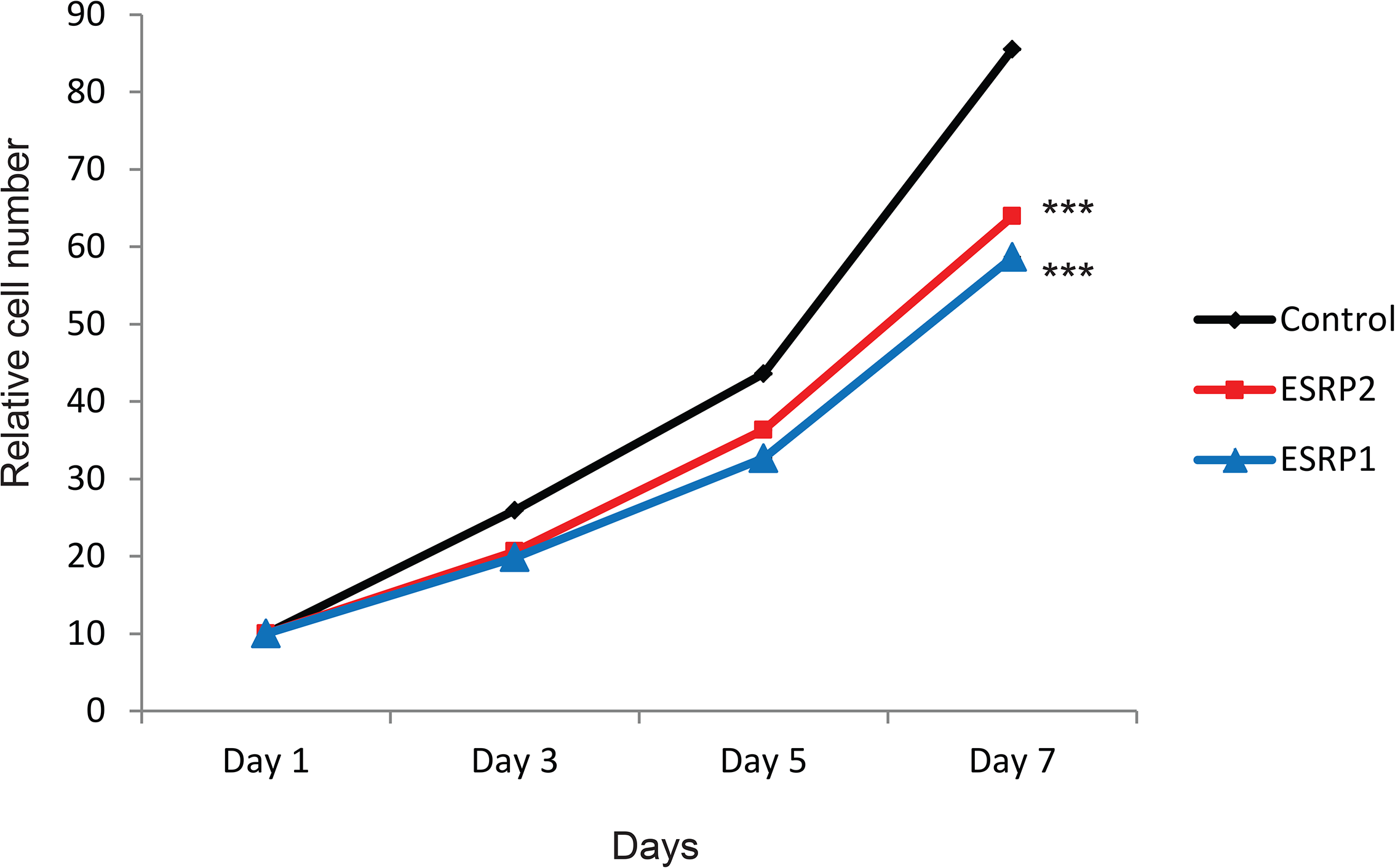
*In vitro* cell proliferation of PC3 and DU145 cells overexpressing ESRP1 or ESRP2 A. Cell growth assay in PC3 cells overexpressing ESRP proteins B. Cell growth assay in DU145 cells overexpressing ESRP proteins

**Figure 5 – Figure Supplement 1.**
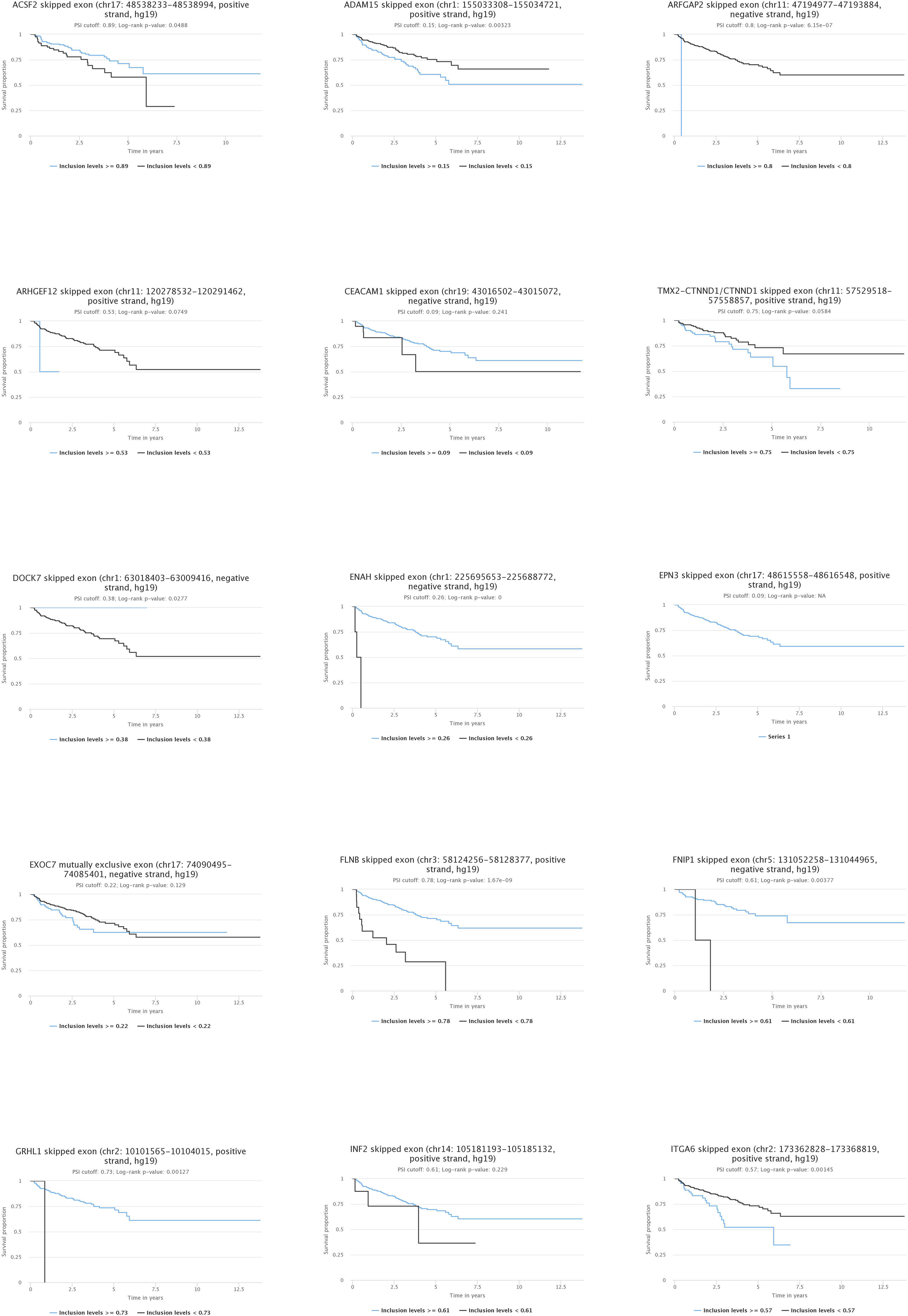
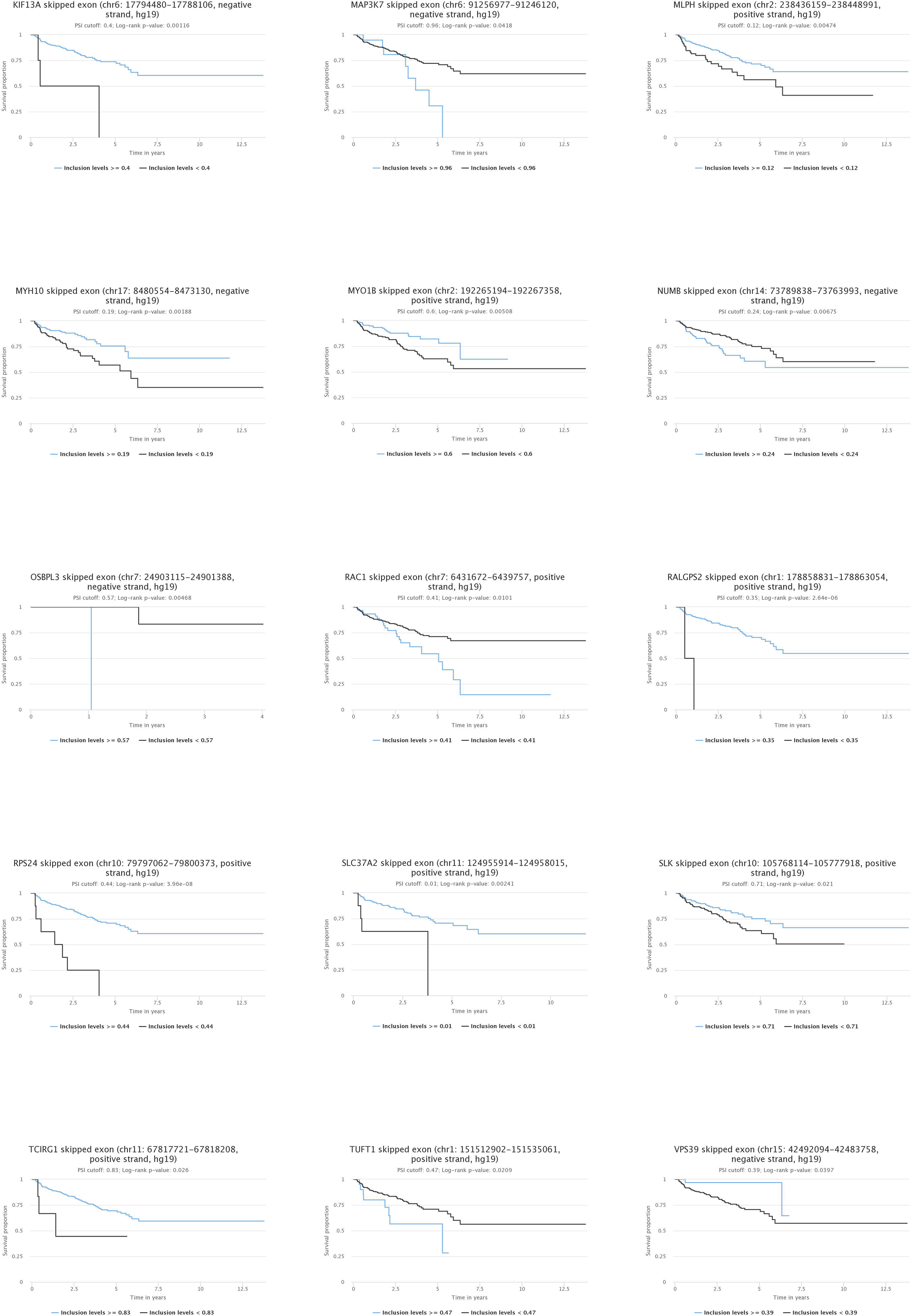
Kaplan-Meier plot showing data from TCGA PRAD cohort of percentage of tumours that are free of biochemical recurrence versus time in years associated with expressing ESRP2-regulated alternative splice isoforms.

**Figure 2 – source data 1.** Meta-analysis of 719 clinical prostate cancer tumours from 11 previously published studies detected significant up-regulation of both *ESRP1* and *ESRP2* in 9/11 datasets

**Figure 3 – source data 1.** Alternative splicing events identified by Suppa2 (55). 446 ESRP regulated alternative splicing events were identified across 319 genes (ΔPSI>10%, p<0.05).

**Figure 3 – source data 2.** Details of 44 experimentally validated ESRP1/ESRP2 target exons identified within prostate cancer cell lines. Gene names (column A) are shown next to PSI levels detected under different experimental conditions (columns B-J); what the pattern of splicing in the PRAD dataset (76) between tumour as compared to normal tissue (Tumour versus normal, column K); the p value associated with the pattern of splicing shown in column K (T-test p-value (BH adjusted), column L); and the difference from the median pattern of inclusion (Δ median PSI, column M) or expression in normal versus prostate tumour tissue in the PRAD cohort (76); whether there was any correlation in the PRAD dataset (76) between splicing inclusion or exclusion of the exon with time to biochemical recurrence of the tumour (column N); the coordinates of the alternative event on hg38 (Alternative event 1 (HG38), column O) and hg19 (Alternative event 1 (HG19), column N); and the forward (column Q) and reverse (column R) primers used to detect the alternative event using RT-PCR.

